# Alzheimer’s disease copathology in dementia with Lewy bodies is associated with astroglial α-synucleinopathy

**DOI:** 10.1101/2024.01.02.573857

**Authors:** Hanne Geut, Emma van den Berg, Baayla D.C. Boon, Jeroen J.M. Hoozemans, Jon-Anders Tunold, Lasse Pihlstrøm, Laura E. Jonkman, J.M. Annemieke Rozemuller, A.W. Evelien Lemstra, Wilma D.J. van de Berg

## Abstract

**Background:** In dementia with Lewy bodies (DLB), co-existence of Alzheimer’s disease (AD) pathology, i.e. amyloid-β plaques and tau tangles, has been associated with a more rapid disease progression. In post-mortem DLB brains, we examined the association between AD copathology and regional load and morphology of α-synuclein pathology. Also, we compared regional load and morphology of AD copathology in DLB to pathology in AD.

**Methods:** We included 50 autopsy-confirmed DLB donors with a clinical DLB phenotype, categorized as having no/low levels of AD copathology (pure DLB, *n* = 15), or intermediate/high levels of AD copathology (mixed DLB+AD, *n* = 35), and autopsy-confirmed pure AD donors (*n* = 14) without α- synuclein pathology. We used percentage area of immunopositivity for quantitative assessment of pathology load, and visual scores for semi-quantitative assessment of different morphologies of α- synuclein, amyloid-β and phosphorylated tau (p-tau) pathology in fifteen neocortical, limbic and brainstem regions.

**Results:** Mixed DLB+AD compared to pure DLB showed a shorter disease duration (6 ± 3 versus 8 ± 3 years, *p* = 0.021) and higher frequency of *APOE*-ε4 alleles. A-synuclein load was higher in neocortical regions (temporal, parietal and occipital), but not in brainstem and limbic regions, which was based upon an increase of Lewy bodies, α-synuclein-positive astrocytes and α-synuclein-positive plaques in these regions. A-synuclein load was most strongly correlated to amyloid-β and p-tau load in temporal (*r* = 0.38 and *r* = 0.50 respectively) and occipital regions (*r* = 0.43 and *r* = 0.42 respectively). Compared to pure AD, mixed DLB+AD showed a lower amyloid-β load in temporal cortex, CA3 and CA4 region, and lower p-tau loads in frontal and parietal cortex, based both upon presence of fewer neuritic plaques as well as neurofibrillary tangles.

**Conclusions:** In DLB brains, AD copathology was associated with more neocortical α-synuclein pathology, consisting not only of Lewy bodies and plaques, but also of astroglial α-synuclein. AD pathology in DLB cases is less than in AD cases, reflecting less advanced pathological stages. Astroglial α-synuclein and its relation with AD copathology in DLB should be further studied, as this may play a role in accelerating clinical decline.

## Background

Dementia with Lewy bodies (DLB) is, after Alzheimer’s disease (AD), the second most common neurodegenerative type of dementia. Next to cognitive decline and dementia, core clinical features of DLB are the presence of cognitive fluctuations, visual hallucinations, rapid eye movement (REM) sleep behaviour disorder (RBD) and parkinsonism (1). Clinical symptoms and disease progression in DLB are heterogeneous, with a median duration from disease onset until death of seven to eight years (2).

The pathological correlate of the clinical DLB syndrome is heterogeneous, but α-synuclein (αSyn) positive Lewy bodies (LBs) and Lewy neurites (LNs) should be present. The current diagnostic criteria for DLB state that more severe Lewy pathology, in combination with less severe or no AD pathology (i.e. neurofibrillary tangles and neuritic plaques), is more likely to explain the presence of a clinical DLB syndrome (1). In addition, the presence of neuronal loss in the substantia nigra (SN) is related to a higher likelihood of the presence of parkinsonism, but this is not a necessary requirement for the DLB pathological diagnosis (1). The presence of LBs and LNs is also characteristic for Parkinson’s disease (PD) and PD with dementia (PDD). As there is much overlap in the neuropathological features of PD, PDD and DLB, the distinction between these entities is primarily defined by the clinical phenotype with an early onset of dementia in the disease course of DLB (1).

In addition to the presence of LBs and LNs, astroglial α-synucleinopathy is frequently observed in limbic and neocortical regions in PD and DLB, and more often in cases with a high load of LBs and LNs (3, 4). Also, up to 50-80% of DLB cases fulfil neuropathological criteria for AD, because of widespread amyloid-β positive plaques, neuritic plaques, and neurofibrillary tangles and threads (5, 6). As such, DLB and AD form a spectrum with four distinct subgroups, namely pure DLB, mixed DLB+AD, AD with amygdala-predominant Lewy pathology and pure AD (6). Besides AD copathology, capillary cerebral amyloid angiopathy (CAA type 1) has been described in 50-90% of DLB cases (7, 8). CAA was found to be present more often in DLB than in PD and PDD (9).

Several clinical studies have demonstrated a more rapid cognitive decline and a higher mortality in clinically probable DLB patients with signs of concomitant AD pathology, assessed with cerebrospinal fluid (CSF) biomarkers (10) and amyloid PET-imaging (11–13). In autopsy-confirmed cohorts, mixed DLB+AD patients also showed a faster cognitive deterioration than pure DLB and pure AD patients (14–16). The combined presence of αSyn and AD pathology may have an additive effect on neurodegeneration and cognitive decline in these patients. However, basic studies have also found evidence for both amyloid-β and tau to act synergistically with αSyn to promote fibril formation and accumulation of each other *in vitro* and *in vivo* (17, 18). This effect may contribute to the accelerated disease course in patients with combined αSyn and AD pathology.

To further broaden our understanding of the relation and synergy between αSyn pathology and AD pathology in DLB, post-mortem examinations studying the regional load as well as morphology of pathological lesions are crucial. To our knowledge, only a few autopsy series have examined the regional load of αSyn, p-tau and amyloid-β pathology in DLB in a quantitative manner using digital pathology (19–22). These and other post-mortem studies identified a positive correlation between AD pathology and the neocortical load of αSyn pathology in DLB (6, 20, 23, 24). A recent study by Coughlin *et al.* compared pure DLB (*n* = 35) to mixed DLB+AD (*n* = 20), and confirmed the higher αSyn burden in neocortical regions in mixed DLB+AD, with an equivalent αSyn burden in entorhinal cortex and putamen (20). However, differences in the morphology of αSyn-positive lesions between pure DLB and mixed DLB+AD patients were not studied. In addition, Coughlin *et al.* found a lower neocortical load of tau pathology in mixed DLB+AD than in pure AD, with a relatively higher tau load in the temporal cortex (20), but the relation of these findings to differences in AD pathological stages remains unclear.

Our aim was to examine whether AD copathology in DLB is related to region-specific differences in α- synuclein load and morphology in a well-characterized autopsy cohort. Based on previous literature (6, 20, 23, 24), we expected an increase in neocortical αSyn pathology in mixed DLB+AD compared to pure DLB. In addition, we tested the hypothesis that regional load and/or morphology of amyloid-β and p-tau pathology over the brain differs between mixed DLB+AD and pure AD, as has been suggested by others (20–22).

## Methods

### Donor selection

We included pure DLB and mixed DLB+AD brain donors, who participated in the brain donation program from the Netherlands Brain Bank (NBB, www.brainbank.nl). For comparison of the distribution of concomitant AD pathology, an age-matched group of fourteen AD brain donors with a clinical and neuropathological diagnosis of AD from the NBB were included in the study (pure AD). For all donors, a written informed consent for brain autopsy and the use of the material and clinical information for research purposes had been obtained from the donor or the next of kin. Criteria for inclusion for both pure DLB and mixed DLB+AD donors in the current study were: 1) presence of limbic- transitional or diffuse-neocortical LB disease (LBD) upon autopsy, and 2) a clinical diagnosis of probable DLB according to the consensus criteria of the DLB Consortium (1). The distinction between the pure DLB group and the mixed DLB+AD group was subsequently made based on the presence and severity of AD copathology. Donors with no or low levels of AD copathology were categorized as ‘pure DLB’. Donors with intermediate or high levels of AD copathology were categorized as ‘mixed DLB+AD’, as at least intermediate levels of AD pathology are considered adequate explanation for cognitive impairment in the absence of other types of pathology (25). Pure AD donors were included based on presence of a typical AD clinical and neuropathological phenotype. Importantly, donors with an atypical clinical AD phenotype, or neuropathological presence of αSyn pathology in brainstem, amygdala, hippocampus or cortex, were excluded. Donors were excluded when insufficient clinical information was available for classification according to these criteria.

Of all donors from the NBB in the period 1989-2015 (*n* = 4003), 439 donors had neuropathological LBD, of whom 259 showed limbic-transitional or neocortical-diffuse LBD upon autopsy. Presence of dementia had been reported in 200 of these donors. Criteria for a clinical diagnosis of ‘probable DLB’ were fulfilled in 52 donors (1). Donors with insufficient available tissue were excluded (*n* = 2). As a result, a group of 15 pure DLB donors and a group of 35 mixed DLB+AD donors were included in the study.

### Clinical information

Demographic features and clinical symptoms were abstracted from the clinical files, including sex, age at symptom onset, age at death, disease duration, presence of dementia, core and supportive clinical features for DLB according to the McKeith criteria (1), and time from disease onset until nursing home placement. At the time of death, the treating physician filled out information on cognitive functioning in the year before death using the Global Deterioration Scale by Reisberg et al. (26) and/or Clinical Dementia Rating (CDR) scale (27), which was converted into the CDR sum of boxes score (CDR-SOB) (28) by adding up the subscores from the different functional domains.

### Neuropathological assessment

Brain regions were dissected according to the standardized procedure of the NBB (open access: www.brainbank.nl). Formalin-fixed paraffin-embedded tissue blocks from the following brain regions were obtained: substantia nigra (SN), amygdala and striatum from the DLB donors, and hippocampus at the level of the lateral geniculate nucleus, middle frontal gyrus, temporal pole, superior parietal lobule and occipital pole from the DLB and AD donors. Seven µm-thick sections were immunostained using antibodies against αSyn (clone KM51, 1:500, Monosan Xtra, The Netherlands), amyloid-β (clone 6F/3D, 1:500, Dako, Denmark for DLB donors, and clone 4G8, 1:8000, Biolegend, USA for AD donors) and hyperphosphorylated tau (p-tau, clone AT8, 1:500, Thermo Fisher Scientific, USA). For all immunostainings, antigen retrieval included incubation in citrate buffer of pH 6.0 for 30 min at 95 ⁰C and subsequent incubation with 80% formic acid for 5 min. Endogenous peroxidase activity was blocked using 1% H2O2 for 15 min, before incubating the sections overnight at 4 ⁰C with the primary antibody in a PBS buffer containing 2% bovine serum albumin. After rinsing in PBS, the sections were incubated with horseradish peroxidase (HRP) secondary mouse antibody (DAKO Envision kit cat# K4001) for 30 min and the staining was visualized using 3,3’-diaminobenzidine (DAB; DAKO cat# K3468). Hippocampus mid and superior frontal gyrus, occipital, superior parietal and medial temporal cortex were immunostained for αSyn, amyloid-β and p-tau. In addition, amygdala was immunostained for αSyn and amyloid-β, and SN was immunostained for αSyn.

For immunofluorescent double-labelling of αSyn inclusions in astrocytes, antigen retrieval was performed as described above. After blocking additional epitopes with 3% normal donkey serum for 30 minutes, sections were incubated overnight at 4 ⁰C with a cocktail of the primary antibodies αSyn (clone KM51 1:500, Monosan Xtra, The Netherlands) and glial fibrillary acidic protein (GFAP, Z0334, 1:4000, DAKO, Denmark). After rinsing in PBS, sections were incubated with secondary antibodies for 2 hr at room temperature (donkey-anti-mouse Alexa 488, 1:200, and donkey-anti-rabbit Alexa 594, 1:200, in blocking buffer containing 2% normal donkey serum and 0.1% Triton-X in TBS (pH 7.6)). Sections were washed in PBS, counterstained using DAPI and coverslipped using Mowiol with 2.5% DABCO.

Braak and McKeith αSyn stages were determined using the BrainNet Europe (BNE) criteria (29). Based on Thal amyloid-β phases scored on the medial temporal lobe (30), Braak neurofibrillary stages (31) and CERAD neuritic plaque scores (32), levels of AD pathology were determined according to NIA-AA consensus criteria (25). For a majority of the pure DLB and mixed DLB+AD cases, we could not make a distinction between Thal phase 4 and 5 as sampling of the cerebellum was not performed. Hence, cases with Thal phase 4 and 5 were analysed as one group. Additionally, Thal CAA stages (33), presence of aging-related tau astrogliopathy (ARTAG) (34), microvascular lesions and hippocampal sclerosis were assessed.

### Quantitative assessment of pathology

All immunostained sections were digitalized using the Vectra Polaris Quantitative Pathology Imaging System (PerkinElmer, USA) at 200x magnification. Regions of interest (ROIs) were manually delineated using ImageJ software (35). Examples of ROIs in different regions are shown in **Supplementary Fig S1**.

All quantitative image analyses were performed in ImageJ using in-house scripts (**Supplementary Methods**). For αSyn load, circular objects (≥ 6 μm in diameter) per cm^2^ were measured in all regions except the SN, to exclude the majority of glial staining, dot-like synaptic staining and neurites from the count (**Fig 1a,b**). For the SN, a different script was used to distinguish between the DAB signal and the neuromelanin pigment. As the selections of LBs were often of irregular shape, object circularity was not used as a criterion for selection in this region (**Fig 1c,d)**. For amyloid-β and p-tau, we measured the percentage area occupied by immunopositivity within each ROI (**Fig 1e-h**). All image selections were visually checked for artefacts, which were manually excluded from the ROIs.

**Figure 1.**
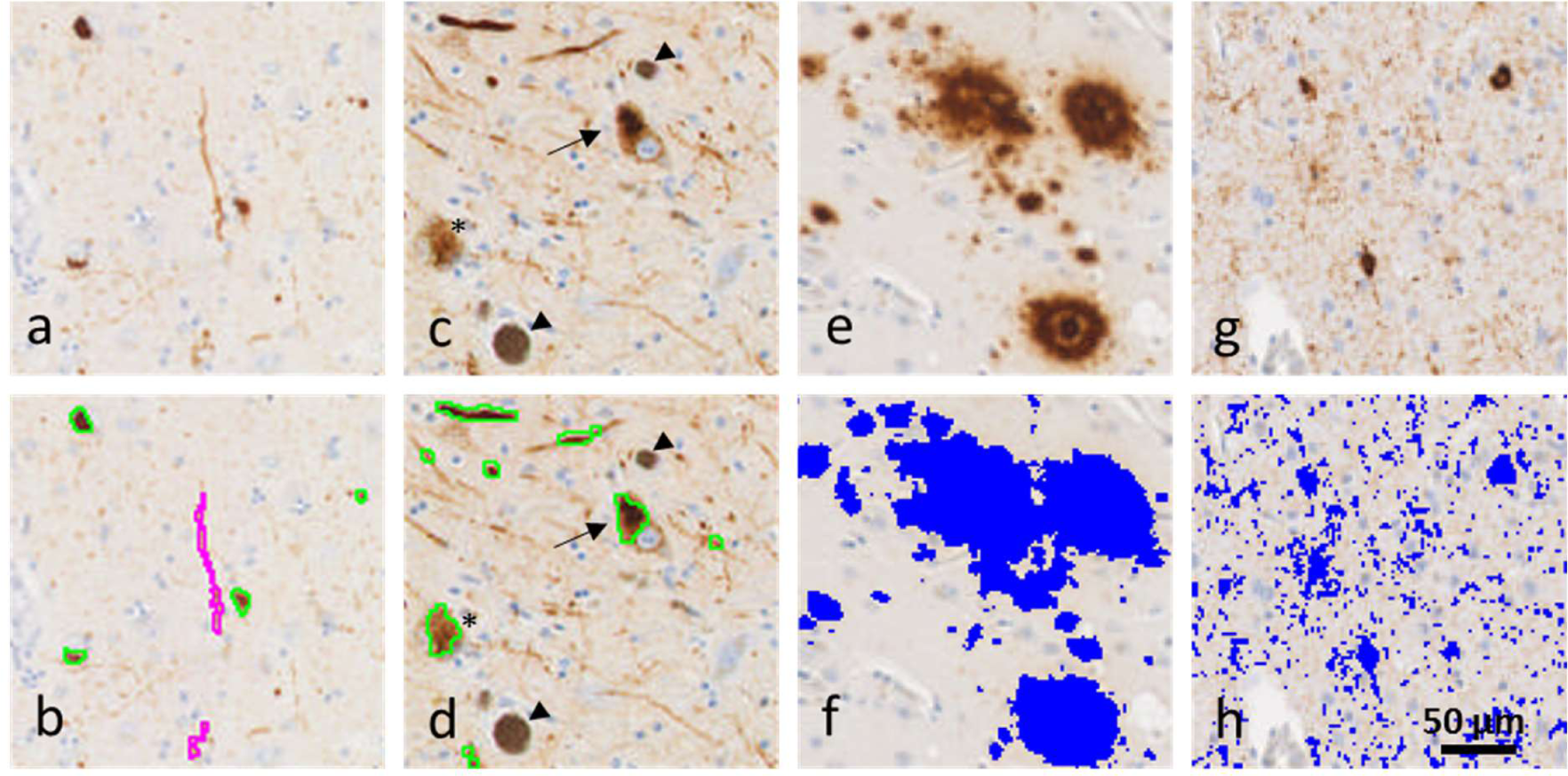
**Selection of pathology using automated ImageJ scripts. a,b**) Circular αSyn positivity, including mainly Lewy bodies (green in **b**) and non-circular αSyn positivity, such as Lewy neurites, plaques and astrocytic pathology (magenta in **b**, data not shown) were selected based on size and circularity of the object using automated image analysis in all regions except for the SN. **c,d**) In the SN, αSyn positivity included Lewy bodies (arrow), granular cytoplasmic intraneuronal immunoreactivity (asterisk) and thick Lewy neurites. Neuromelanin was distinguished from DAB based on hue and saturation of the color (arrowheads). **e-h**) Percent surface area of immunopositivity was measured for amyloid-β (**e,f**) and p-tau (**g,h**). The scale bar in h applies to all images.

### Qualitative assessment of pathology

We compared qualitative scores of αSyn-positive morphological structures between pure DLB and mixed DLB+AD. We assessed the following αSyn-positive morphologies in the SN, amygdala, CA2, CA1, parahippocampal gyrus, medial temporal, superior frontal, superior parietal and occipital cortex: LBs, granular cytoplasmic neuronal immunoreactivity (GCNIR), long LNs, dots and short neurites, astrocytic positivity and globular plaques (36) (**Fig 3a-h**). We visually rated the morphologies on an ordinal scale ranging from 0 to 3 (**Supplementary Table S1**). In the SN, halo-like LBs and compact LBs were scored separately (29). LBs, LNs, GCNIR and dots have been described in the BNE consensus criteria (29). The presence of αSyn in the dystrophic neurites of amyloid-β plaques is well-known (37–39). This can be visible as a patch of globular structures, sometimes surrounding an empty core (globular plaques, **Fig 3h**). Astrocytic αSyn-positivity may be present as coiled body-like inclusions or as star-like astrocytes (3, 36, 38, 40, 41). A gallery of αSyn-positive astrocytic lesions in different brain regions of the DLB donors is shown in **Supplementary Fig S2a-i**. Double labelling experiments showed αSyn-positive inclusions within the cell soma of astrocytes, visualized using GFAP, in various brain regions of DLB donors (**Supplementary Fig S2j-o**).

Additionally, the morphology of amyloid-β-positive pathology was assessed and compared between the three study groups. We distinguished eight different amyloid-β positive morphological structures.

Seven of these types have been described in the BNE criteria: classic plaques, compact plaques, diffuse plaques, subpial band-like positivity, pial positivity, cerebral amyloid angiopathy (CAA) type 1 (capillary and dyshoric angiopathy) and CAA type 2 (42) (**Fig 5a-c,e-h**). Furthermore, amyloid-β-positive coarse-grained plaques were assessed, a recently described type of plaque associated with inflammatory and vascular changes (43) (**Fig 5d**). All types of morphology were assessed in the temporal and frontal cortex, except for CAA type 1 and 2 which were assessed in the occipital and frontal cortex (**Supplementary Table S1**). Ordinal scores ranging from 0 to 3 were averaged over the two regions.

Finally, we compared p-tau positive morphologies between the three study groups. We distinguished between three types of AD-related morphological structures, i.e. neuropil threads, neurofibrillary tangles and neuritic plaques (31) (**Fig 5j-l**), and two types of age-related tau astrogliopathy (ARTAG), namely thorn-shaped astrocytes and fuzzy astrocytes (34) (**Fig 5m,n**). P-tau pathology was assessed in the parahippocampal cortex and temporal pole according to a well-defined ordinal scoring system (**Supplementary Table S1**). Averages of the ordinal scores have been reported.

### *APOE* genotyping

*APOE* alleles were determined by genotyping of rs429358 and rs7412 as part of a larger sample set using either the TaqMan® SNP Genotyping Assay (ThermoFisher Scientific) for 13 of the AD cases, or the Infinium® NeuroChip Consortium Array v1.1 (Illumina) (44) for 46 of the DLB cases. GenomeStudio 2.0 was used to call variants. Additional quality checks were performed on the *APOE* data from the DLB cases, using plink 1.9 and included standard filtering of variants and individuals based on missingness, Hardy-Weinberg equillibrium, relatedness, excess heterozygosity, sex-check and ancestry assessed by principal components. Two mixed DLB+AD cases had an ambiguous *APOE* ε1/ε3 or ε2/ε4 genotype, which was interpreted as ε2/ε4 in further analyses.

### Statistical analysis

To examine between-group differences regarding clinical and pathological features, we used independent sample *t*-tests for continuous variables, chi-square tests for categorical variables and Mann-Whitney U tests for ordinal variables.

Distribution tests showed a right-skewed and leptokurtic distribution for αSyn load (skewness 3.5 [standard error 0.09] and kurtosis 17.5 [SE 0.18]) and amyloid-β load (skewness 1.7 [SE 0.08] and kurtosis 3.7 [SE 0.16]) with relatively normal distributed data for p-tau pathology (skewness 1.0 [SE 0.09], kurtosis -0.3 [SE 0.17]). A log-transformation (*y* = ln(*x* + 1)) was performed on the data from the three types of pathology to normalize the data distribution (log-αSyn load: skewness -0.7, kurtosis 0.5; log-amyloid-b skewness 0.1, kurtosis -1.1; log-p-tau skewness -0.1, kurtosis -1.5). Linear mixed effect models were used to compare the log-transformed quantitative pathological load and the visual scores between groups and across brain regions. We included group, region and group*region as fixed factors, and age at death as covariate in the model. Post-hoc tests were performed to test differences between the groups within regions, using Bonferroni-correction for the number of tested regions in each analysis. As the Thal phases and Braak neurofibrillary stages were higher in pure AD than in mixed DLB+AD, we performed subgroup analyses where only cases with Thal phase 4 or 5 were assessed for differences in amyloid-β load and only cases with Braak NFT phase 5 or 6 were assessed for differences in p-tau load. Data analysis was performed using IBM SPSS Statistics version 26. A significance level of *p* < 0.05 was maintained for all tests after Bonferroni-correction for multiple testing.

Hierarchical clustering of αSyn-positive morphologies, DLB donors based on regional αSyn-positive morphologies and all donors based on log-transformed regional amyloid-β and p-tau loads and neocortical amyloid-β and p-tau morphologies was performed using the *pheatmap* package version 1.0.12 in R version 3.6.1 using default settings.

## Results

### Shorter survival in mixed DLB+AD compared to pure DLB and pure AD

Demographics and main clinical features of the 15 pure DLB donors, 35 mixed DLB+AD donors and 14 pure AD donors are presented in **Table 1**. The percentage of males was similar between groups (80%, 66% and 71% in pure DLB, mixed DLB+AD and pure AD respectively). Age at symptom onset and age at death were not significantly different between groups. However, disease duration was shorter in the mixed DLB+AD group compared to the pure DLB group (6 ± 3 vs. 8 ± 3 years, *p* = 0.021) and compared to the pure AD group (10 ± 5 years, *p* = 0.019).

**Table 1.** Clinical and genetic characteristics of the study groups.

|  | Pure DLB | Mixed DLB+AD | Pure AD | P-value^<br>Pure DLB vs.<br>mixed DLB+AD | P-value^<br>Mixed DLB+AD<br>vs. pure AD |
| --- | --- | --- | --- | --- | --- |
| **N** | 15 | 35 | 14 |  |  |
| **Sex M/F (%M)** | 12/3 (80%) | 23/12 (66%) | 10/4 (71%) | .3 | .7 |
| **Age at death y**<br>mean ± SD (range) | 75 ± 8 (61-90) | 75 ± 8 (56-87) | 76 ± 12 (53-92) | .8 | .8 |
| **Age at onset y**<br>mean ± SD (range) | 67 ± 7 (53-79) | 69 ± 8 (50-83) | 66 ± 10 (48-84) | .5 | .2 |
| **Disease duration y**<br>mean ± SD (range) | 8 ± 3 (3-13) | 6 ± 3 (2-14) | 10 ± 5 (2-23) | .021 | .019 |
| **Post-mortem delay hh:mm**,<br>mean ± SD (range) | 4:59 ± 0:52<br>(4:00-6:34) | 5:00 ± 1:06 (2:15-7:52) | 7:15 ± 2:05<br>(3:05-11:33) | 1.0 | .001 |
| **APOE genotype** |  |  |  |  |  |
| N (%) | 13 | 33 | 13 | .000 | .2 |
| ε4 non-carrier | 8 (62%) | 3 (9%) | 4 (31%) | | |
| ε4 heterozygous | 5 (39%) | 26 (79%) | 8 (62%) | | |
| ε4 homozygous | 0 (0%) | 4 (12%) | 1 (8%) | | |

^ Groups were compared using t-tests for continuous variables, chi-square tests for categorical variables and Mann-Whitney U test for ordinal scores (i.e. APOE genotype). APOE = apolipoprotein E.

The severity of dementia in the year before death, as measured by the CDR-SOB (28) or the Reisberg scale (26), was not different between the pure DLB and mixed DLB+AD group. Although not reaching significance, we observed higher frequencies of depressive episodes (63% vs. 33%, *p* = 0.055, **Table 2**) and an earlier admission to a nursing home (4 ± 2 vs. 6 ± 3 years, *p* = 0.059) in mixed DLB+AD compared to pure DLB.

**Table 2.**
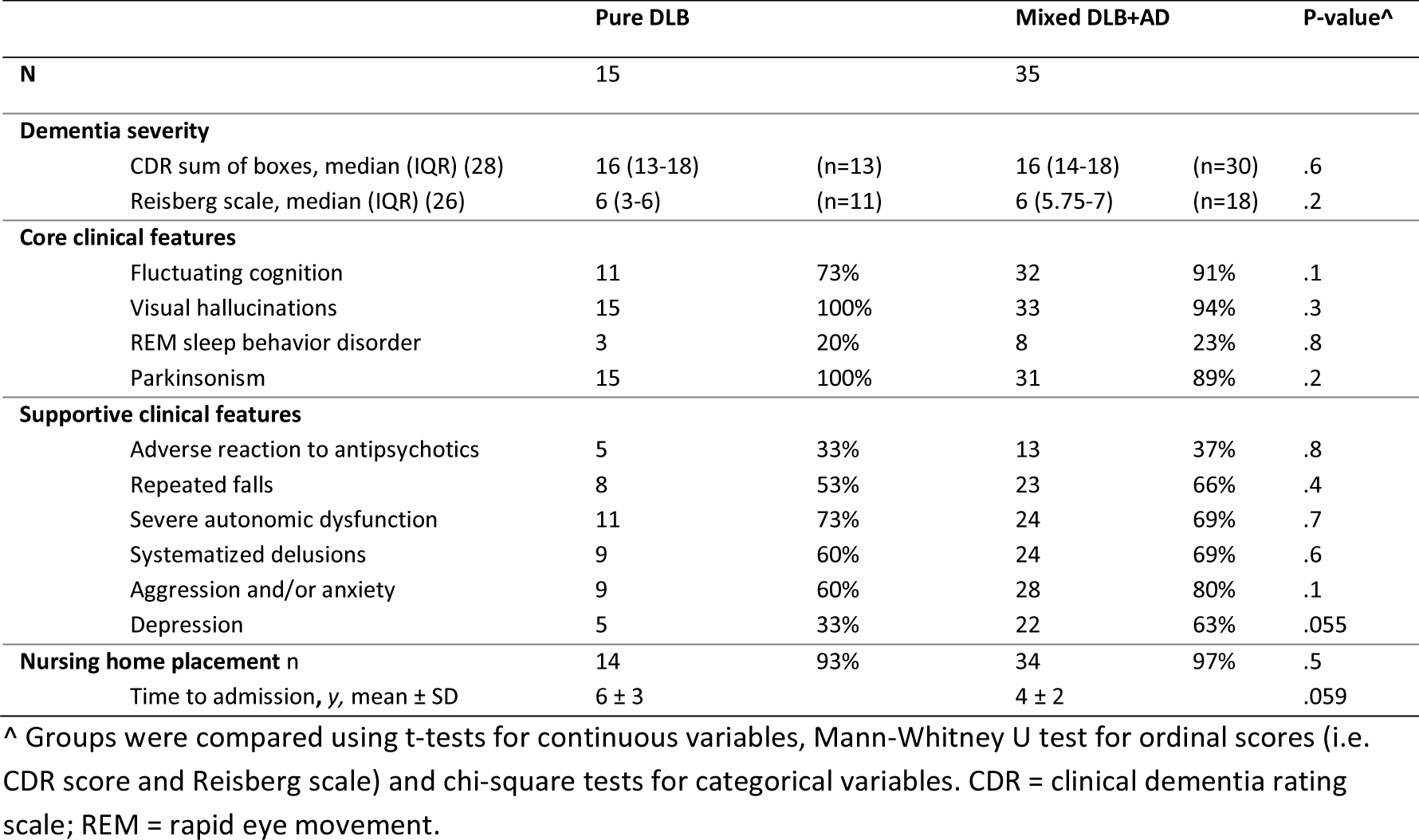
Clinical symptoms according to McKeith criteria. (**45**) **in pure DLB and mixed DLB+AD.**

### Mixed DLB+AD showed less advanced AD pathology and less frequent CAA type 1 than pure AD

The neuropathological features and stages have been summarized in **Table 3**. As by definition, the mixed DLB+AD group showed higher Thal amyloid-β phases, Braak neurofibrillary stages and CERAD scores, and higher levels of AD pathology (all *p* < 0.001) than the pure DLB group. In pure AD, Thal amyloid-β phases were marginally higher than in mixed DLB+AD, but this difference did not reach significance (*p* = 0.1). Braak neurofibrillary stage and CERAD scores were higher in pure AD than in mixed DLB+AD (*p* = 0.001 and *p* < 0.001 respectively). Braak and McKeith LB stages were not significantly different between the two DLB groups. Also, presence of ARTAG, microvascular lesions and hippocampal sclerosis was not significantly different between the three study groups. CAA type 1, the capillary form of CAA, was more common in the mixed DLB+AD donors compared to pure DLB (49% vs. 7%, *p* = 0.005), with six cases reaching stage 2 or 3 (17%). CAA type 1 was highly frequent in the pure AD group (79%), with 3 cases reaching stage 2. CAA type 2 was equally present in pure DLB (47%), mixed DLB+AD (51%) and pure AD (21%).

**Table 3.**
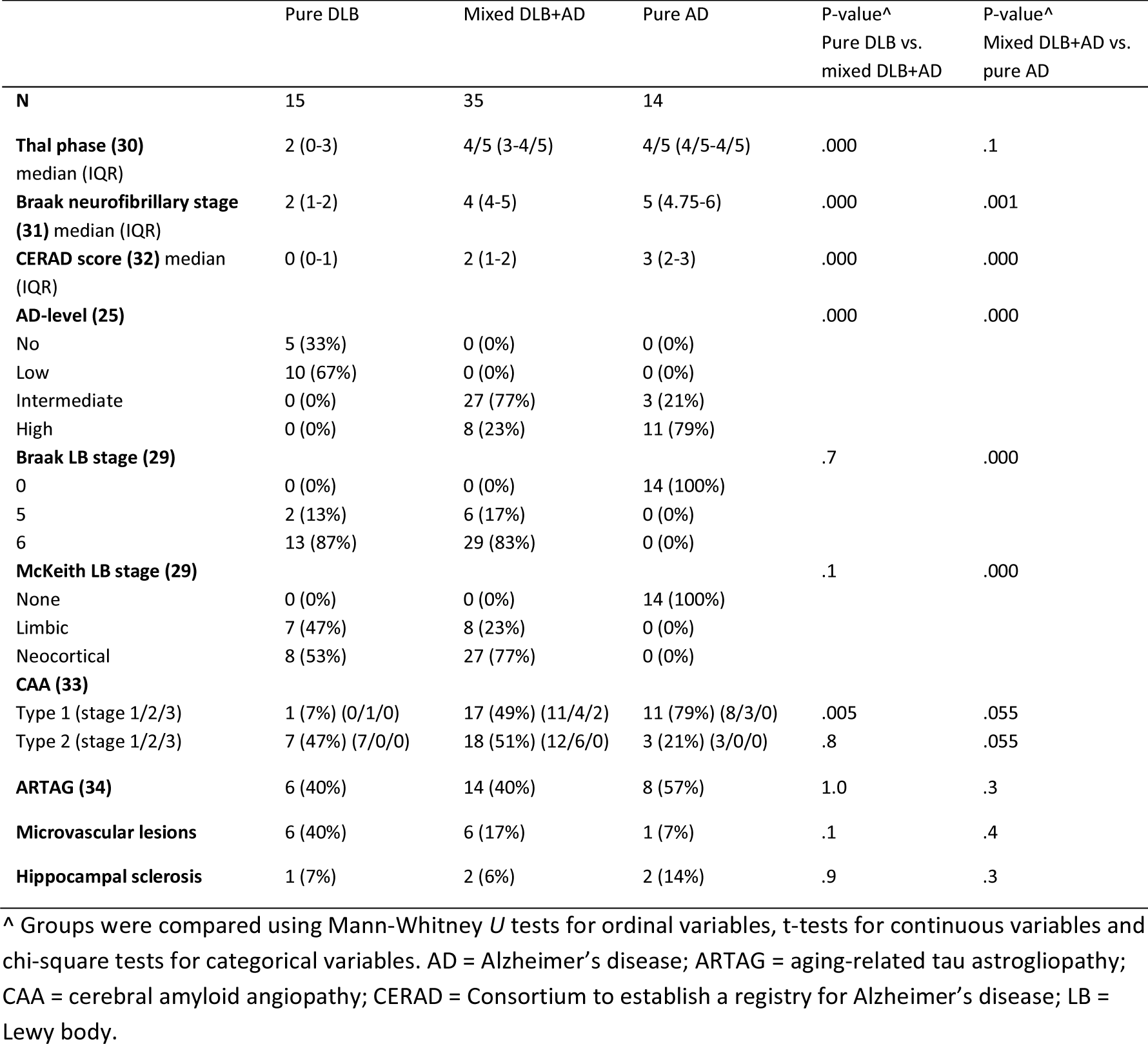
Neuropathological features and stages of the study groups.

|  | Pure DLB | Mixed DLB+AD | Pure AD | P-value^<br>Pure DLB vs.<br>mixed DLB+AD | P-value^<br>Mixed DLB+AD vs.<br>pure AD |
| --- | --- | --- | --- | --- | --- |
| **N** | 15 | 35 | 14 |  |  |
| **Thal phase (30)**<br>median (IQR) | 2 (0-3) | 4/5 (3-4/5) | 4/5 (4/5-4/5) | .000 | .1 |
| **Braak neurofibrillary stage (31)** median (IQR) | 2 (1-2) | 4 (4-5) | 5 (4.75-6) | .000 | .001 |
| **CERAD score (32)** median (IQR) | 0 (0-1) | 2 (1-2) | 3 (2-3) | .000 | .000 |
| **AD-level (25)** |  |  |  | .000 | .000 |
| No | 5 (33%) | 0 (0%) | 0 (0%) |  |  |
| Low | 10 (67%) | 0 (0%) | 0 (0%) |  |  |
| Intermediate | 0 (0%) | 27 (77%) | 3 (21%) |  |  |
| High | 0 (0%) | 8 (23%) | 11 (79%) |  |  |
| **Braak LB stage (29)** |  |  |  | .7 | .000 |
| 0 | 0 (0%) | 0 (0%) | 14 (100%) |  |  |
| 5 | 2 (13%) | 6 (17%) | 0 (0%) |  |  |
| 6 | 13 (87%) | 29 (83%) | 0 (0%) |  |  |
| **McKeith LB stage (29)** |  |  |  | .1 | .000 |
| None | 0 (0%) | 0 (0%) | 14 (100%) |  |  |
| Limbic | 7 (47%) | 8 (23%) | 0 (0%) |  |  |
| Neocortical | 8 (53%) | 27 (77%) | 0 (0%) |  |  |
| **CAA (33)** |  |  |  |  |  |
| Type 1 (stage 1/2/3) | 1 (7%) (0/1/0) | 17 (49%) (11/4/2) | 11 (79%) (8/3/0) | .005 | .055 |
| Type 2 (stage 1/2/3) | 7 (47%) (7/0/0) | 18 (51%) (12/6/0) | 3 (21%) (3/0/0) | .8 | .055 |
| **ARTAG (34)** | 6 (40%) | 14 (40%) | 8 (57%) | 1.0 | .3 |
| **Microvascular lesions** | 6 (40%) | 6 (17%) | 1 (7%) | .1 | .4 |
| **Hippocampal sclerosis** | 1 (7%) | 2 (6%) | 2 (14%) | .9 | .3 |

^ Groups were compared using Mann-Whitney *U* tests for ordinal variables, t-tests for continuous variables and chi-square tests for categorical variables. AD = Alzheimer's disease; ARTAG = aging-related tau astrogliopathy; CAA = cerebral amyloid angiopathy; CERAD = Consortium to establish a registry for Alzheimer's disease; LB = Lewy body.

### αSyn load was increased in neocortical regions of mixed DLB+AD compared to pure DLB

We compared αSyn load between mixed DLB+AD and pure DLB in SN, limbic and neocortical regions. Although αSyn load in SN, hippocampal subregions, amygdala and frontal cortex was similar between the DLB groups, αSyn load was increased in the medial temporal gyrus (*p* = 0.002), superior parietal (*p* = 0.006) and occipital cortex (*p* = 0.006) in mixed DLB+AD compared to pure DLB (**Fig 2a**).

**Figure 2.**
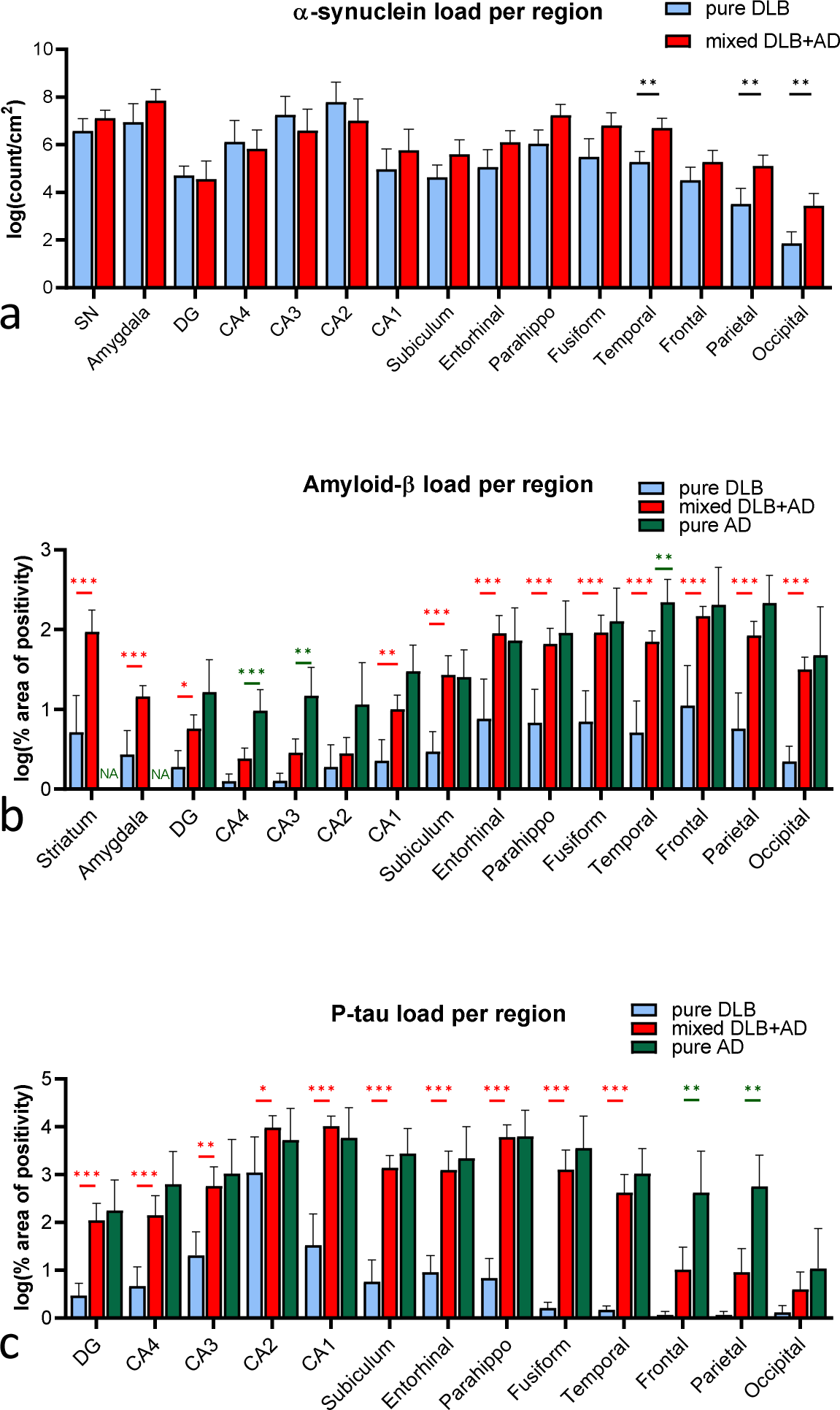
Alpha-synuclein, amyloid-β and p-tau load showed region-specific differences between disease groups. a) Alpha-synuclein load a was increased in mixed DLB+AD compared to pure DLB in the temporal, parietal and occipital cortex. **b)** Amyloid-β load was higher in pure AD compared to mixed DLB+AD in the temporal cortex, CA3 and CA4 regions. **c)** P-tau load was higher in mixed DLB+AD than in pure DLB in all brain regions except for the frontal, parietal and occipital cortex. P-tau load was higher in pure AD than in mixed DLB+AD in the frontal and parietal cortex. * *p* < 0.05, ** *p* < 0.01, *** *p* < 0.001.

### Region-specific increase in αSyn-positive morphologies in mixed DLB+AD compared to pure DLB

We studied differences in the presence of the observed αSyn-positive morphologies (**Fig 3a-h**) between the two DLB groups. Astrocytic coiled-body-like inclusions were only observed in the SN, whereas star-like astrocytes as well as globular plaques were not observed in the SN, but were present in all other regions. Mixed DLB+AD compared to pure DLB showed a significant increase in the global presence of all types of αSyn-positive morphologies except for coiled body-like inclusions (LBs, GCNIR, dots, plaques, star-like astrocytes: all p < 0.001; LNs: p = 0.002; coiled body-like inclusions: p = 0.7).

Post-hoc comparisons between regions showed a significant increase in LBs in the parahippocampal and medial temporal cortex (*p* = 0.009 and *p* < 0.001), in dots and short neurites in the temporal cortex (*p* = 0.038), in star-like astrocytes in the parahippocampal (*p* = 0.002) and temporal cortex (*p* = 0.003) and in globular plaques in the CA1 region (*p* = 0.028), parahippocampal cortex (*p* = 0.007) and temporal cortex (*p* < 0.001, **Fig 3i, Supplementary Fig S3a-g**).

Data-driven analysis, using hierarchical clustering based on the regional semi-quantitative load of the previously defined αSyn positive morphologies, showed clustering of all pure DLB donors and a few mixed DLB+AD donors in cluster 1 (*n* = 15, **Supplementary Fig S3h**), with mild to moderate scores for LBs, LNs and αSyn positive dots, and no or little astrocytic pathology and αSyn positive plaques. Cluster 2 (*n* = 10) consisted of mixed DLB+AD donors with mild to severe LB, LN and dot-like pathology, but little or no astrocytic pathology and plaques. Cluster 3 (n = 7) and cluster 4 (n = 9) showed moderate to severe LB, LN and dot-like pathology, combined with severe plaque pathology in parahippocampal gyrus and temporal cortex (cluster 3), and/or severe astrocytic αSyn pathology (cluster 4).

### Lower amyloid-β load in temporal cortex and hippocampal subregions in mixed DLB+AD than in pure AD

Amyloid-β pathology and p-tau pathology loads were assessed using surface area measurements of immunopositivity and compared between the three study groups. As expected, amyloid-β load was increased in mixed DLB+AD compared to pure DLB in all assessed regions except for the CA2, CA3 and CA4 regions (DG: *p* = 0.028; CA1: *p* = 0.002; striatum, amygdala, subiculum, and entorhinal, parahippocampal, fusiform, temporal, frontal, parietal and occipital regions: all *p* < 0.001, **Fig 2b**). When comparing mixed DLB+AD to pure AD, amyloid-β load was lower in the medial temporal cortex (*p* = 0.009), CA3 region (*p* = 0.002) and CA4 region (*p* > 0.001) in mixed DLB+AD (**Fig 2b**). To examine the effect of the more advanced level of AD pathology in the AD cases, we performed a subanalysis on only the cases with Thal amyloid-β phase 4 or 5 (mixed DLB+AD: *n* = 21; pure AD: *n* = 9). Here, similar regional differences in amyloid-β load between mixed DLB+AD and pure AD were observed (lower in in the temporal cortex (*p* = 0.014), CA1 region (*p* = 0.040), CA3 region (*p* < 0.001) and CA4 region (*p* = 0.007) in mixed DLB+AD).

In addition, as expected, p-tau load was increased in mixed DLB+AD compared to pure DLB in all hippocampal and parahippocampal subregions (CA3: *p* = 0.001; CA2: *p* = 0.035; DG, CA4, CA1, subiculum, entorhinal cortex, parahippocampal cortex and fusiform gyrus: all *p* < 0.001), and temporal cortex (*p* < 0.001) (**Fig 2c**). Cluster analysis of all cases based on regional load of amyloid-β and p-tau showed a specific subgroup of pure DLB donors with isolated high levels of p-tau in the CA2 region, while p-tau levels in other regions and amyloid-β levels in all regions were low in these donors (**Supplementary Fig S4**).

P-tau load was lower in mixed DLB+AD than in pure AD in the frontal (*p* = 0.003) and parietal cortex (*p* = 0.002, **Fig 2c**). To determine whether this difference can be explained by the lower Braak NFT stages in mixed DLB+AD compared to pure AD, we analysed only DLB cases and AD cases with Braak neurofibrillary stage 5 or 6 (mixed DLB+AD: *n* = 11; pure AD: *n* = 9). In this comparison, the differences in p-tau load in the frontal and parietal cortex did not reach significance. This suggests that the differences in regional p-tau load between mixed DLB+AD and pure AD are largely based upon the more advanced Braak neurofibrillary stages in pure AD cases.

### AD pathology is most strongly related to αSyn load in the medial temporal and occipital cortex

To study the relationship between αSyn, amyloid-β and p-tau pathology, we examined the region-specific correlation between load of the three pathologies within the two DLB groups (**Fig 4**). The amyloid-β load and p-tau load were significantly correlated in all regions except for CA2, most strongly in the temporal cortex (Pearson’s *r* = 0.63, *p* < 0.001). Load of neocortical αSyn was significantly related to amyloid-β pathology in the parietal (*r* = 0.36, *p* = 0.011), temporal (*r* = 0.38, *p* = 0.010), occipital (*r* = 0.43, *p* = 0.002) and entorhinal cortex (*r* = 0.31, *p* = 0.036). p-Tau load was significantly related to αSyn load in the temporal cortex (*r* = 0.050, *p* = 0.001), occipital cortex (*r* = 0.42, *p* = 0.002), fusiform gyrus (*r* = 0.41, *p* = 0.007), parahippocampal cortex (*r* = 0.42, *p* = 0.003) and subiculum (*r* = 0.29, *p* = 0.047). The temporal and occipital cortex showed the strongest relation between αSyn load and load of amyloid-β and p-tau.

**Figure 3.**
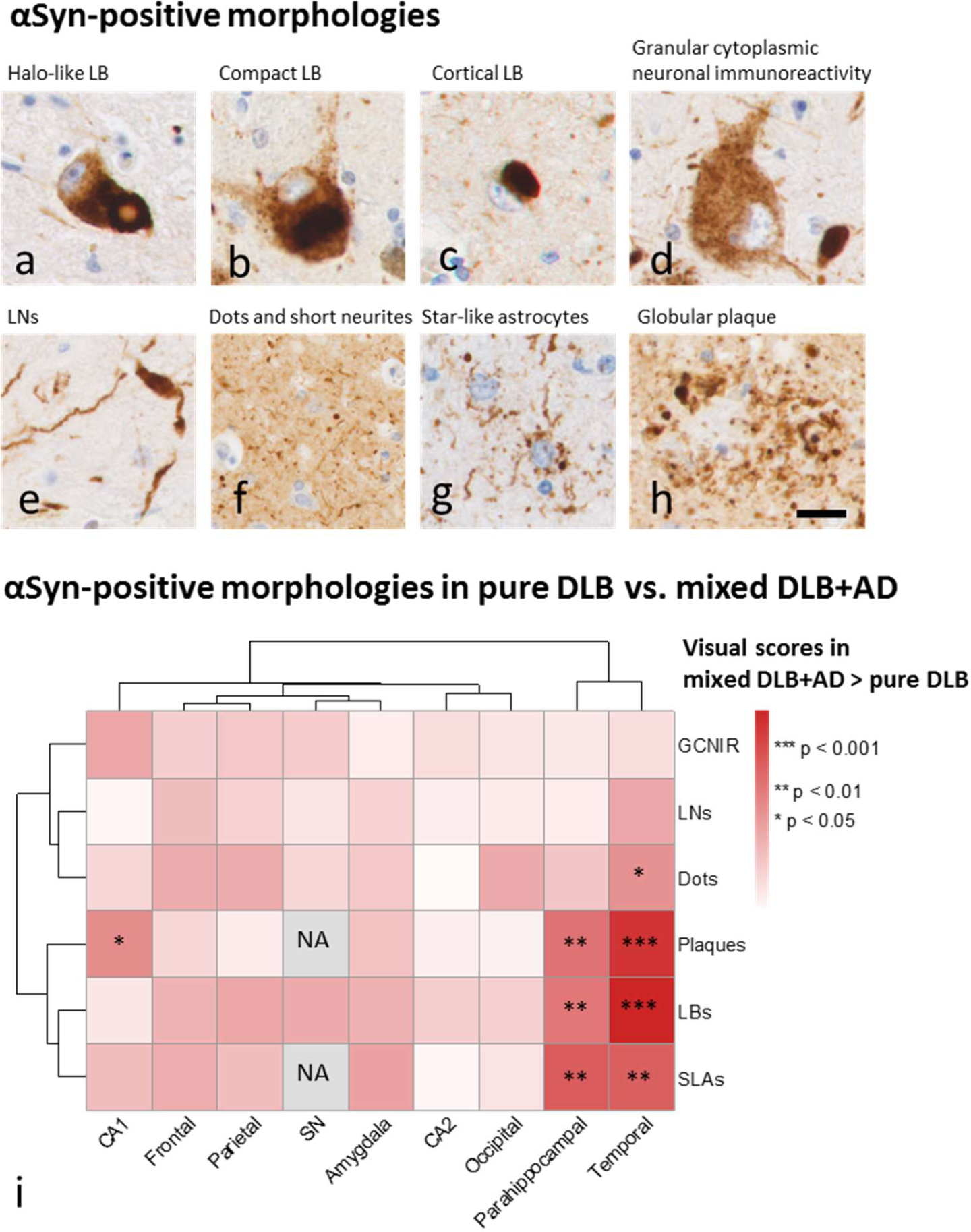
Morphology of αSyn-positivity in pure DLB compared to mixed DLB+AD. Different types of neuronal, glial and extracellular morphology (**a-h**) were assessed in brainstem, limbic and neocortical brain regions. In the SN, we distinguished halo-like (**a**) and compact LBs (**b**). In the other regions, the load of cortical LBs was assessed (**c**). Other types of morphology included **d**) granular cytoplasmic neuronal immunoreactivity (GCNIR), **e**) long LNs, **f**) dots and short neurites, **g**) star-like astrocytes and **h**) globular plaques. **i**) The visual scores for different morphological types of α-synuclein pathology were compared between mixed DLB+AD and pure DLB. Mixed DLB+AD showed higher scores than pure DLB for all types of pathology. Colors represent Bonferroni-corrected – log10 p-values of the post-hoc comparisons per region, using linear mixed modeling while controlling for age. LBs, star-like astrocytes and plaques in the parahippocampal and temporal cortex, dots in the temporal cortex and plaques in the CA1 region were significantly higher in mixed DLB+AD than in pure DLB. The scale bar in h represents 20 μm and applies to panels **a-h**. GCNIR = granular cytoplasmic neuronal immunoreactivity; LBs = Lewy bodies; LNs = Lewy neurites; SN = substantia nigra; SLAs = star-like astrocytes.

**Figure 4.**
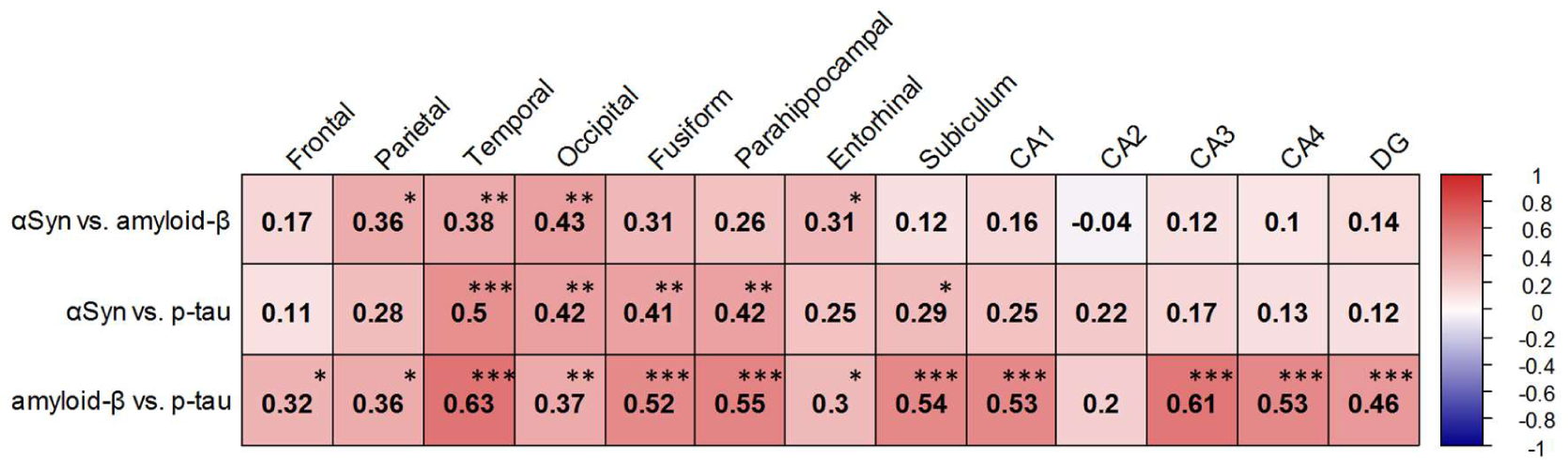
Pearson’s correlation between αSyn, amyloid-β and P-tau load in the two DLB groups. P-tau and amyloid-β load are most strongly related to αSyn load in the temporal and occipital cortex. * *p* < 0.05, ** *p* < 0.01, *** *p* < 0.001.

**Figure 5.**
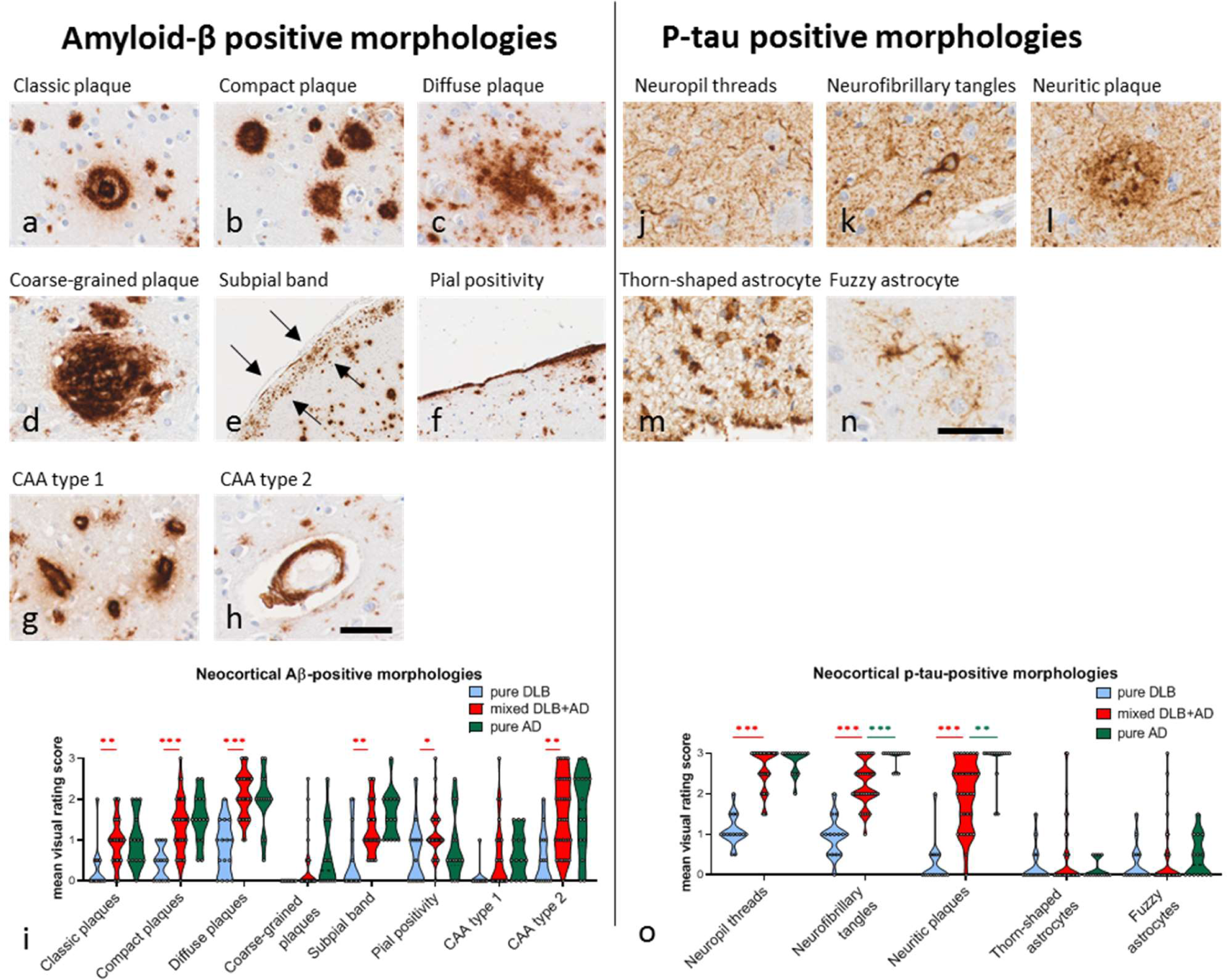
Qualitative assessment of various morphologies of amyloid-β and p-tau pathology. We scored the following types of amyloid-β positive morphologies: **a**) classic plaques, **b**) compact plaques, **c**) diffuse plaques, **d**) coarse-grained plaques, **e**) subpial band-like positivity (between arrows), **f**) pial positivity, **g**) cerebral amyloid angiopathy (CAA) type 1 (i.e. capillary CAA and dyshoric angiopathy), and **h**) CAA type 2. Scale bar in **h** represents 50 μm in **a-d**, **g** and **h**, 500 μm in **e** and 100 μm in **f**. **i**) Burden of classic, compact and diffuse plaques, subpial band-like positivity and CAA type 2 were increased in neocortical regions of mixed DLB+AD compared to pure DLB. Aβ morphologies were not significantly different between mixed DLB+AD and pure AD. Morphological p- tau positive structures included **j**) neuropil threads, **k**) neurofibrillary tangles, **l**) neuritic plaques, **m**) thorn-shaped astrocytes and **n**) fuzzy astrocytes. Scale bar in n represents 50 μm in **j-n**. Images taken in the temporal cortex (**j- l,n**) and subpial layer in hippocampus (**m**). **o**) AD-related p-tau morphologies (threads, tangles and plaques) were more often observed in mixed DLB+AD than in pure DLB, and neurofibrillary tangles and neuritic plaques were more often observed in pure AD than in mixed DLB+AD. Age-related tau pathology of the astroglia (ARTAG, i.e. thorn-shaped astrocytes and fuzzy astrocytes) was observed equally in all groups. CAA = cerebral amyloid angiopathy.

### Similar amyloid-β and p-tau morphologies were present in mixed DLB+AD and pure AD

To compare the morphology of AD pathology between mixed DLB+AD and pure AD, we visually assessed different morphological structures of amyloid-β pathology (**Fig 5a-h**) and p-tau positive pathology (**Fig 5j-n**) in neocortical regions. As by definition, mixed DLB+AD compared to pure DLB showed a higher neocortical burden of classic, compact and diffuse plaques, subpial band-like positivity and CAA type 2 (**Fig 5i**). Coarse-grained plaques were frequently observed in mixed DLB+AD (26%) and pure AD cases (50%), but not in pure DLB. Also, by definition, AD-related p-tau positive morphologies, i.e. neuropil threads, neurofibrillary tangles and neuritic plaques, were all more often present in mixed DLB+AD than in pure DLB. Interestingly, ARTAG-related morphologies, i.e. thorn-shaped astrocytes and fuzzy astrocytes, were equally present in both DLB groups (**Fig 5o**).

When comparing mixed DLB+AD to pure AD, the morphology of amyloid-β-positive lesions was similar (**Fig 5i**). However, p-tau-positive neurofibrillary tangles and neuritic plaques were more frequent in pure AD than in mixed DLB+AD, reflecting the higher Braak NFT stages in pure AD (**Fig 5o**).

We performed hierarchical clustering of all donors with a complete dataset based on the visual scores of the neocortical amyloid-β and p-tau morphologies (**Supplementary Fig S5**). In Cluster 1, 13 out of 15 pure DLB donors were separated from the mixed DLB+AD. Mixed DLB+AD and pure AD donors did not separate based on the morphology of AD pathology.

### *APOE* ε4 alleles were related to neocortical amyloid-β load, but not αSyn or p-tau load

The frequency of *APOE* ε4 alleles was significantly higher in mixed DLB+AD (79% heterozygous, 12% homozygous ε4 carrier) compared to pure DLB (39% heterozygous, 0% homozygous; *p* < 0.001), and similar when compared to pure AD (62% heterozygous, 8% homozygous; p = 0.2, **Table 1**). We tested the correlation between the number of *APOE* ε4 alleles and pathology loads in neocortical regions within the 50 DLB cases. There was a significant correlation between *APOE* ε4 and the load of amyloid- β in frontal cortex (Spearman’s *r* = 0.52, *p* < 0.001), parietal cortex (*r* = 0.50, *p* < 0.001), temporal cortex (*r* = 0.54, *p* < 0.001) and occipital cortex (*r* = 0.60, *p* <0.001). In contrast, the load of αSyn and p-tau in these regions did not significantly correlate to the number of *APOE* ε4 alleles.

## Discussion

In the current study, we aimed to examine whether AD copathology in DLB is associated with region-specific differences in αSyn load and morphology. Mixed DLB+AD showed a higher load of neocortical αSyn than pure DLB, with more astroglial αSyn aggregates. In addition, we compared amyloid-β and p-tau load and morphology between DLB and AD. Load of AD pathology showed region-specific differences between mixed DLB+AD and pure AD, but the morphology was similar.

Similar to results from previous studies (5, 6, 10, 23), the mixed DLB+AD donors in our study showed a shorter disease duration than the pure DLB donors. The shorter survival in clinical DLB than in clinical AD cases is well known from clinical follow-up studies (46). The shorter survival combined with a similar dementia severity at time of death, suggests a faster clinical decline in mixed DLB+AD than in pure DLB. This is further supported by a higher rate of aggression, anxiety and depression, and an earlier nursing home placement in mixed DLB+AD. This is in line with a recent study by Ferman *et al*., who showed an association between more rapid cognitive decline and a more widespread distribution of tau and aSyn pathology in DLB cases (16). This suggests that pathological differences between DLB patients, for example the presence of concomitant AD pathology, burden of neocortical αSyn and presence of astroglial αSyn, contribute to clinical heterogeneity and prognosis in DLB. Besides a potential additive effect of multiple pathologies on cognitive decline, evidence from *in vitro* and *in vivo* studies indicates that both amyloid-β and tau have a synergistic relation with αSyn to enhance each other’s aggregation and accumulation (17, 18). Both additive as well as synergistic effects of these copathologies may therefore play a role in the faster disease progression of mixed DLB+AD cases.

A higher neocortical αSyn load in mixed DLB+AD compared to pure DLB is known from the literature (19, 20, 47), and confirmed by our study. We showed that the increase in αSyn pathology in mixed DLB+AD does not only consist of an increase in LBs and αSyn positivity in dystrophic neurites of senile plaques (observed as globular plaques in the αSyn staining) (37–39), but also of an increase in αSyn- positive astrocytes in the parahippocampal and temporal cortex. Notably, astroglial αSyn in other neocortical regions was exclusively observed in the mixed DLB+AD group, which may represent more advanced stages of astroglial α-synucleinopathy. These observations suggest that astroglial αSyn may contribute to the more rapid clinical decline in mixed DLB+AD compared to pure DLB.

Presence of astroglial α-synucleinopathy has previously been described in PD and DLB, and has been related to the burden of LBs and LNs (3, 48). Interestingly, astrocytes only express very low levels of αSyn under normal conditions, and αSyn-immunopositive astrocytes have also been observed in regions that do not develop Lewy pathology, such as the striatum and dorsal thalamus (3). The mechanism by which astrocytes accumulate αSyn aggregates, is currently under debate. Proposed mechanisms include the removal of toxic extracellular αSyn by phagocytosis (49) or direct transfer of toxic αSyn from neurons via exosomes or nanotubules leading to a prion-like transmission of αSyn pathology (50, 51). Our results indicate that there may be a relation between the accumulation of astroglial αSyn and the presence of AD pathology. This correlation could be mediated by the high abundance of LBs in cases with high levels of AD pathology, as astrocytes may take up neuronal αSyn via scavenging or via direct uptake (3, 48, 50, 51). In addition, an increased neuro-inflammatory response may exacerbate the astrocytic uptake of neuronal αSyn, as was demonstrated in a transgenic mouse model expressing human αSyn (52). Increased microglial activation is a well-known phenomenon in AD (53), and has been described to accompany congophilic plaques in mixed DLB+AD (54). As to whether this astrocytic response is neuroprotective or damaging, and also whether astrocytic uptake of αSyn plays a role in the putative synergistic relationship between αSyn and AD pathology, remains a matter of speculation. Further studies into the mechanisms underlying the relation between astroglial α-synucleinopathy and AD pathology are warranted.

The largest differences in amyloid-β load between pure DLB and mixed DLB+AD were observed in cortical regions, striatum and amygdala, but not the CA3 and CA4 regions of the hippocampus, whereas the largest differences in p-tau load were observed in the CA regions of the hippocampus and temporal cortex, but not the frontal, parietal and temporal cortex. These are the regions that are affected in early pathological stages (**Fig 6a**). When comparing mixed DLB+AD to pure AD, load of amyloid-β and p-tau was similar in early affected regions, and lower in regions affected in more advanced pathological stages (**Fig 6b**). These findings can largely be attributed to the more advanced stages of AD pathology in pure AD. This is corroborated by the subanalysis in cases with advanced Braak NFT stages, where regional differences in p-tau load between groups disappeared. Although this effect was not observed for regional differences in amyloid-β load in a similar subanalysis of cases with advanced amyloid-β Thal phases, as the regional differences between mixed DLB+AD and pure AD remained present in this subgroup, this should be interpreted with caution. The group sizes in this subanalysis are small, and the largest differences were still observed in regions affected in late disease stages. Also, we could not discriminate between Thal phase 4 and 5 in most of the cases, because the cerebellum had not been sampled. Therefore, the pure AD cases in this subgroup may still have a more advanced amyloid-β pathological stage than the mixed DLB+AD cases, explaining the higher pathology load in these late affected regions. These findings on amyloid-β and p-tau load in mixed DLB+AD vs. pure AD diverge from the findings by Coughlin et al., who mainly found a relatively higher p-tau load in the temporal cortex of mixed DLB+AD (20, 22). This may be due to differences in case selection (i.e. PDD and DLB are pooled as one group in the studies of Coughlin et al., and we have excluded AD cases with α- synucleinopathy in the amygdala in our study), as well as methodological differences in use of antibodies, ROI drawing of hippocampal subfields, scripts for image analysis, and statistical analyses (including the use of relative vs. absolute differences). Larger studies in well-defined cohorts are needed to resolve these controversies.

**Figure 6.**
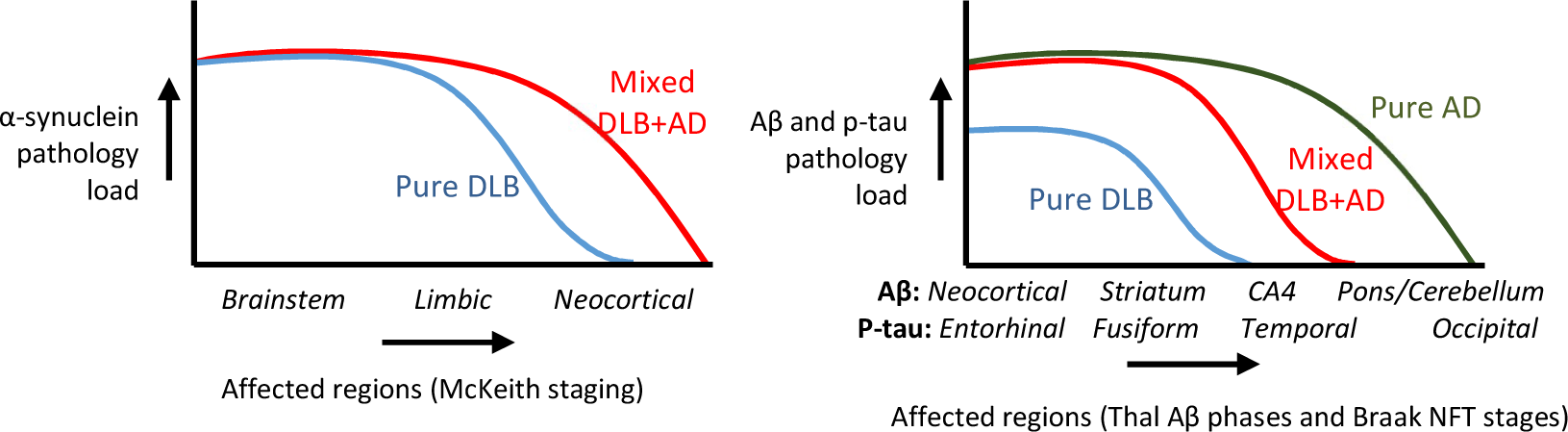
**Schematic representation of regional αSyn, amyloid-β and p-tau pathology in the three study groups. a**) αSyn load is significantly higher in neocortical regions in mixed DLB+AD compared to pure DLB. **b**) Regions affected in earlier Thal amyloid-β phases and Braak NFT stages showed the largest differences in amyloid-β and p-tau load between the DLB groups, whereas regions affected in more advanced pathology stages showed the largest difference between mixed DLB+AD and pure AD.

The presence of amyloid-β and p-tau-positive morphologies in mixed DLB+AD and pure AD was largely similar, except for a higher abundance of neurofibrillary tangles and threads in pure AD. Again, this difference may be based on the higher Braak neurofibrillary stages in pure AD. The almost complete absence of CAA type 1 in pure DLB compared to the frequent presence of CAA type 1 in mixed DLB+AD is in line with the well-known association between CAA type 1 and both Thal amyloid-β phases and Braak neurofibrillary stages (33, 55).

The *APOE* ε4 locus has been reported as an important genetic risk factor for DLB (56, 57). The *APOE* ε4 allele was more frequent in mixed DLB+AD compared to pure DLB. This is in line with the findings from other autopsy series of PDD and DLB cases (23, 58, 59) and with clinical studies in DLB (10). Recently, several studies have found evidence for a direct relation between the *APOE* ε4 locus and αSyn pathology, independent from amyloid-β pathology, both in mouse models (60, 61), as well as in post-mortem brains from LBD patients (60, 62). However, we did not find any evidence for an independent association between the *APOE* ε4 locus and neocortical load of αSyn pathology. The differences in the findings from the two post-mortem studies may be due to different inclusion criteria, as both studies included brains from PD and DLB cases, whereas our study included only DLB cases. Therefore, there are only few cases in our study without a high load of neocortical αSyn pathology. Together with the small sample size, our study may lack statistical power to detect an association between APOE ε4 and neocortical αSyn pathology. Second, the association found in the other studies may also be driven by the inclusion of PD cases, and may lack within the DLB subgroup. In the study by Dickson et al., the association was strongest within cases with none or little AD pathology, a group that will predominantly consist of PD cases. Further studies of larger DLB autopsy series are needed to elucidate this matter.

This is one of the largest autopsy series of DLB cases including both digital pathology as well as qualitative scoring to determine the load and morphology of pathological structures. The selection of DLB cases based on the clinical probable DLB phenotype, and the separation from cases with a PDD phenotype, enabled us to study the clinicopathological correlations within this clinical entity. Limitations of the current study include the small sample size of the pure DLB and pure AD group. Future studies may include other types of concomitant pathologies, such as TDP-43 pathology, as well as inflammatory markers.

## Conclusions

In conclusion, a higher neocortical α-synuclein load, consisting of LBs, astroglial α-synucleinopathy and plaques, was related to AD copathology in DLB. Morphology of AD copathology in mixed DLB+AD was similar to pathology in pure AD, but the load was lower in regions affected in advanced pathological stages. This suggests that besides the presence of AD copathology, these neocortical αSyn-positive morphologies, in particular astroglial α-synucleinopathy, may play a role in the more rapid clinical decline in mixed DLB+AD compared to pure DLB. Subtyping DLB based on presence of AD copathology may provide insight into pathogenic mechanisms contributing to disease progression in DLB, and may guide the selection of patients for novel therapeutic strategies targeting αSyn and amyloid-β pathology.

## List of abbreviations

αSyn: alpha-synuclein
AD: Alzheimer’s disease
ARTAG: aging-related tau astrogliopathy
CA: cornu ammonis
CAA: cerebral amyloid angiopathy
CDR: clinical dementia rating scale
CDR-SOB: clinical dementia rating scale – sum of boxes score
CGP: coarse-grained plaque
CSF: cerebrospinal fluid
DG: dentate gyrus
DLB: dementia with Lewy bodies
GCNIR: granular cytoplasmic neuronal immunoreactivity
p-tau: phosphorylated tau
LB: Lewy body
LBD: Lewy body disease
LN: Lewy neurite
NBB: Netherlands Brain Bank
PD: Parkinson’s disease
PDD: Parkinson’s disease with dementia
RBD: rapid eye movement (REM) sleep behaviour disorder
ROI: region of interest
SN: substantia nigra

## Declarations

### Ethics approval and consent to participate

All procedures of the NBB donation program have been approved by the medical ethics committee of the VU University Medical Center (Amsterdam UMC, location VUmc, Amsterdam).

### Consent for publication

For all donors, a written informed consent for brain autopsy and the use of the material and clinical information for research purposes had been obtained from the donor or the next of kin.

### Availability of data and materials

The data that support the findings of this study are available from the corresponding author upon reasonable request and with permission of the Netherlands Brain Bank. The data are not publicly available due to privacy or ethical restrictions

### Authors’ contributions

HG and EvdB selected the DLB donors, and designed, and undertook pathological experiments of the DLB donors. BB and LEJ selected the AD donors, and designed, and undertook pathological experiments of the AD donors. HG and EvdB performed all qualitative and quantitative image analyses. JAT and LP provided and interpreted APOE data. JMR provided a pathological diagnosis of all donors and supervised the morphological assessment of pathological structures. AWL retrospectively diagnosed DLB donors according to clinical criteria. HG interpreted data and wrote the initial draft of the manuscript. AWL and WDJvdB provided overall study direction, supervision, and revised the manuscript. All authors critically reviewed the report and approved the final version.

## Acknowledgements

The authors thank John Bol and Irene Frigerio for excellent technical assistance.

