## Supplementary materials for "Alzheimer’s disease copathology in dementia with Lewy bodies is associated with astroglial α-synucleinopathy"

### Supplementary information

#### Supplementary Methods

##### ImageJ macro for circular and non-circular αSyn load

Pathological load of αSyn was defined as the count of immunopositive objects per $cm^2$. The LBs were separated from other immunopositive structures based on size and circularity, to exclude αSyn-positive plaques, dots, astrocytic pathology and LNs. In short, the DAB channel was selected with the Color Deconvolution plugin [1]. Then, diffuse synaptic staining and background staining was subtracted using Gaussian blurring. After thresholding with the MaxEntropy autothresholding algorithm, circular objects (circularity 0.5-1) with a surface area of at least 36 $\mu m^2$ (i.e. largest diameter of ≥6 µm) were selected, as this identified most LBs but not most extracellular dots (example in green in **Fig 1a,b**). Non-circular objects (circularity 0-0.5) with a surface area of at least 16 $\mu m^2$ were measured separately (example in magenta in **Fig 1a,b**, data not shown).

##### ImageJ macro for αSyn load in the SN

In the SN, a different macro was used to distinguish DAB-positivity from neuromelanin pigment (**Fig 1c,d**). The DAB signal was selected from the raw image using HSB color thresholding, with the same parameters for all images. Pixels were merged using a dilation/erosion step, and particles with a size of at least 50 $\mu m^2$ (i.e. ≥ 7x7 to 1x50 µm) were counted.

##### ImageJ macro for amyloid-β and p-tau load

The amyloid-β and p-tau load were defined as percentage surface area of DAB-positivity in the scanned regions (examples in **Fig 1e-h**). The DAB channel was selected using the Color Deconvolution plugin. The DAB image was thresholded using the MaxEntropy autothresholding algorithm, and the percentage area was measured within each ROI.

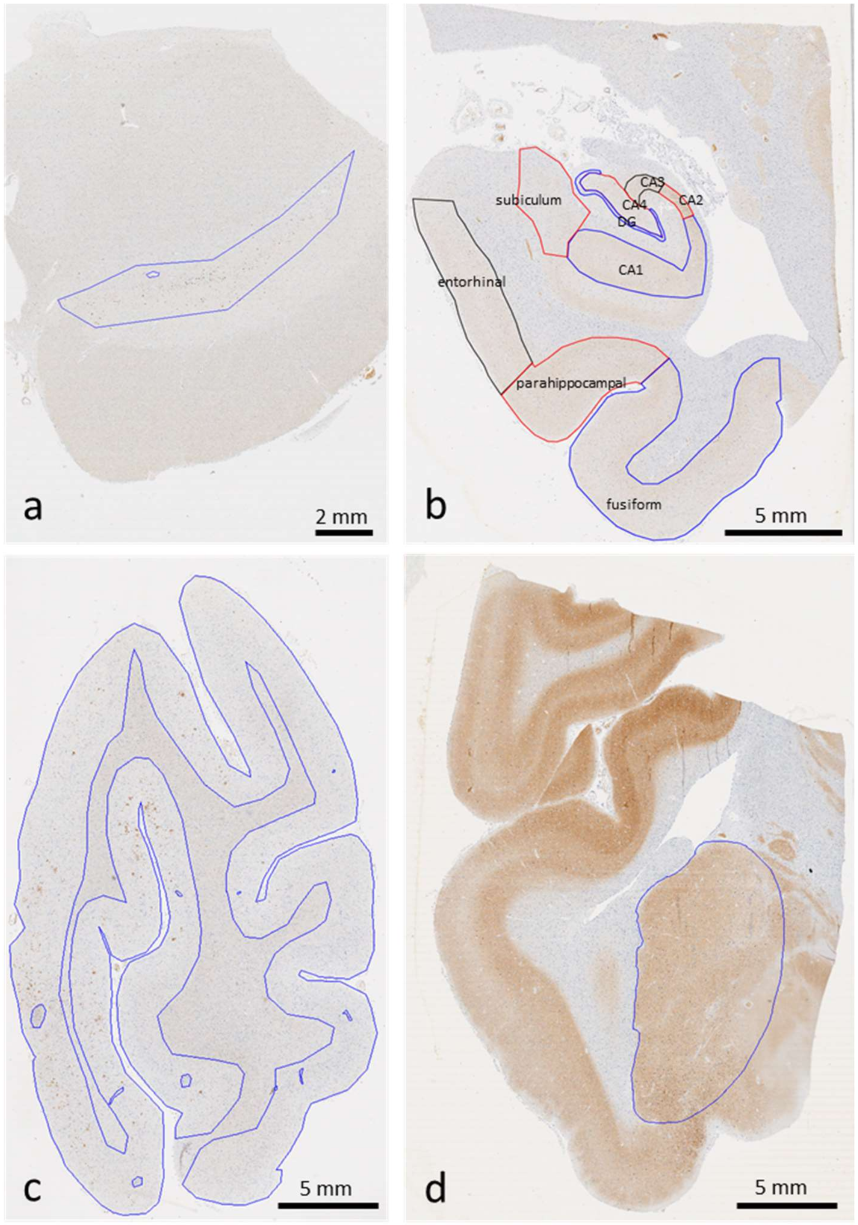


**Supplementary Figure S1. Examples of ROIs in various brain regions.** **a**) Substantia nigra pars compacta and pars reticulata were included. **b**) Hippocampal ROIs included dentate gyrus (DG), cornu ammonis (CA) regions 1-4, subiculum, entorhinal cortex, parahippocampal gyrus and fusiform gyrus, whenever present in the section. **c**) Cortical grey matter cut perpendicular to the white matter was included for all neocortical regions. In the occipital cortex, both striate and peristriate gray matter was included. **d**) The entire amygdaloid complex was included.

**Supplementary Table S1.** Criteria used for visual assessment of αSyn, amyloid-β and p-tau positive structures.

| | Regions scored | Ordinal scoring | | | |
|---|---|---|---|---|---|
| | | **1** | **2** | **3** | |
| **Alpha-synuclein immunoreactive structures** | | | | | |
| Halo-like Lewy body<br>Compact Lewy body<br>Granular cytoplasmic neuronal immunoreactivity | Substantia nigra | ≤1 | 2-4 | >4 | out of 10 neurons |
| Coiled body-like inclusions | Substantia nigra | 1-3 | 4-10 | >10 | in the entire region |
| Cortical Lewy body<br>Star-like astrocytes<br>Globular plaques | Limbic and cortical regions | 0-1 | 2-5 | >5 | in a 20x field |
| Lewy neurites | All regions | 0-1 | 2-5 | >5 | in a 20x field |
| Dots and small neurites | All regions | just visible at 10x field | easily visible at 10x field, empty patches of neuropil in between | macroscopically visible, neuropil densely covered | |
| **Amyloid-β immunoreactive structures** | | | | | |
| Classic plaque<br>Compact plaque<br>Diffuse plaque | Temporal and frontal cortex | 0-1 | 2-5 | >5 | in a 20x field |
| Coarse-grained plaque | Temporal and frontal cortex | 1-3 | >3, solitary | >3, in groups | in the entire region |
| Subpial band<br>Pial positivity | Temporal and frontal cortex | individual plaques / short lengths of positivity | band-like, long uninterrupted lengths, <50% of the pial lining | band-like / pial positivity in >50% of the pial lining | |
| CAA type 1 | Occipital and frontal cortex | 1-5 | >5, solitary | >20, in groups | positive capillaries in entire region |
| CAA type 2 | Occipital and frontal cortex | only in meningeal vessels | <50% of cortical vessels | >50% of cortical vessels | positive vessels in entire region |
| **P-tau immunoreactive structures** | | | | | |
| Neuropil threads | Hippocampus and temporal cx | just visible at 10x field | easily visible at 10x field | macroscopically visible, neuropil densely covered | |
| Neurofibrillary tangles | Hippocampus and temporal cx | 1-5 | 6-20 | >20 | in a 20x field |
| Neuritic plaques | Hippocampus and temporal cx | 0-1 | 2-5 | >5 | in a 20x field |
| Thorn-shaped astrocytes | Hippocampus and temporal cx | occasional clusters | frequent clusters | numerous/ widespread | in entire region |
| Fuzzy astrocytes | Hippocampus and temporal cx | 1-2 | 3-10 | >10 | in a 20x field |

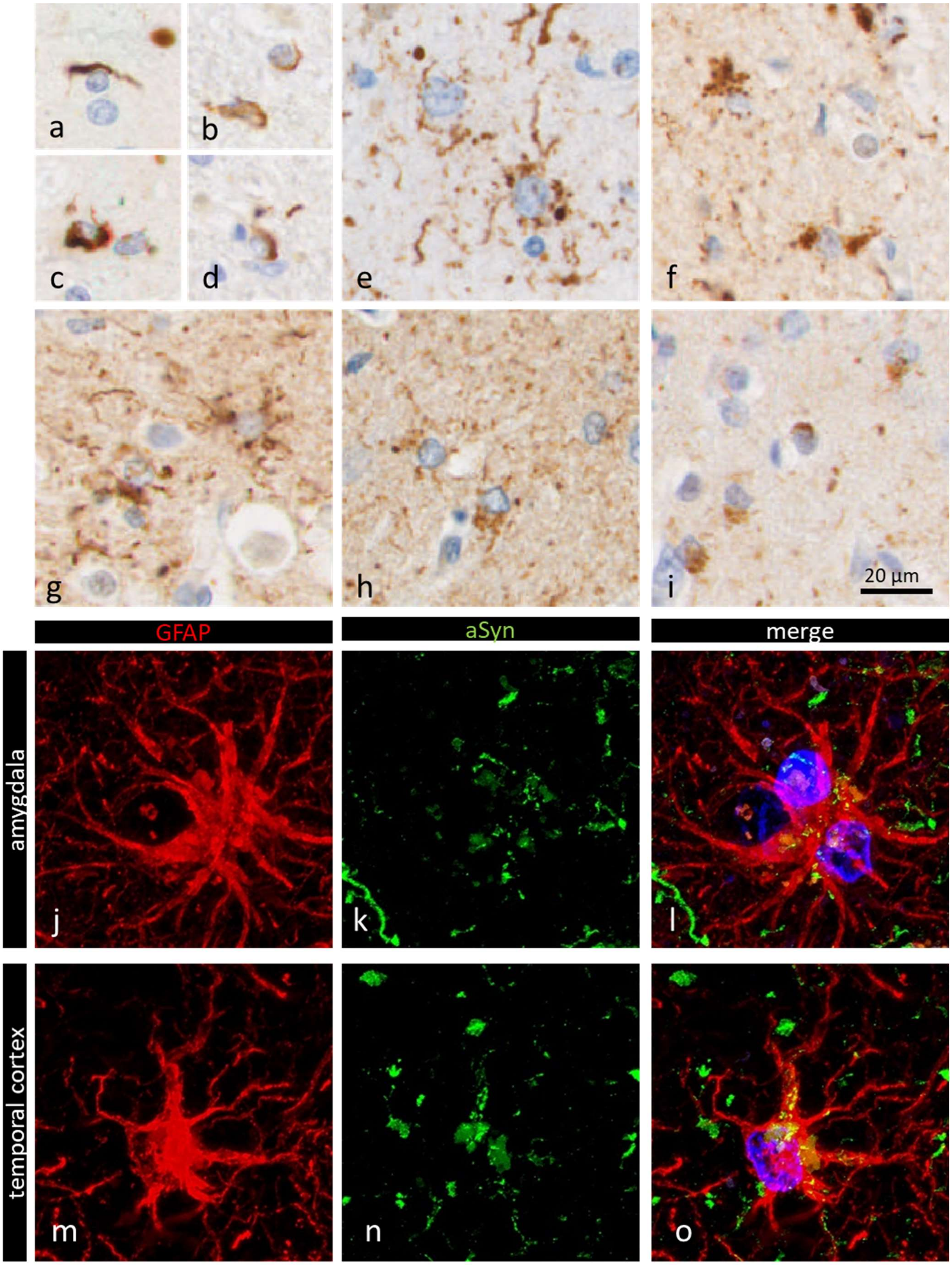
a
b
c
d
e
f
g
h
i
20 µm
GFAP
aSyn
merge
amygdala
temporal cortex
j
k
l
m
n
o

**Supplementary Figure S2. Astrocytic αSyn-positive inclusions in various brain regions of DLB donors**, using antibody clone KM51. **a-d**) Examples of coiled-body like inclusions in the midbrain. **e-i**) Different appearances of star-like astrocytes (SLA) in **e**) the amygdala, **f**) the CA1 region, **g**) the parahippocampal gyrus, **h**) the temporal cortex, and **i**) the frontal cortex. Astrocytes (GFAP, red) with αSyn-positive inclusions (clone KM51, green) in the amygdala (**j-l**) and temporal cortex (**m-o**). Representative images were taken in four mixed DLB+AD donors with a high load of astroglial α-synucleinopathy. The scale bar in **i** applies to images **a-i**.

### Supplementary Results

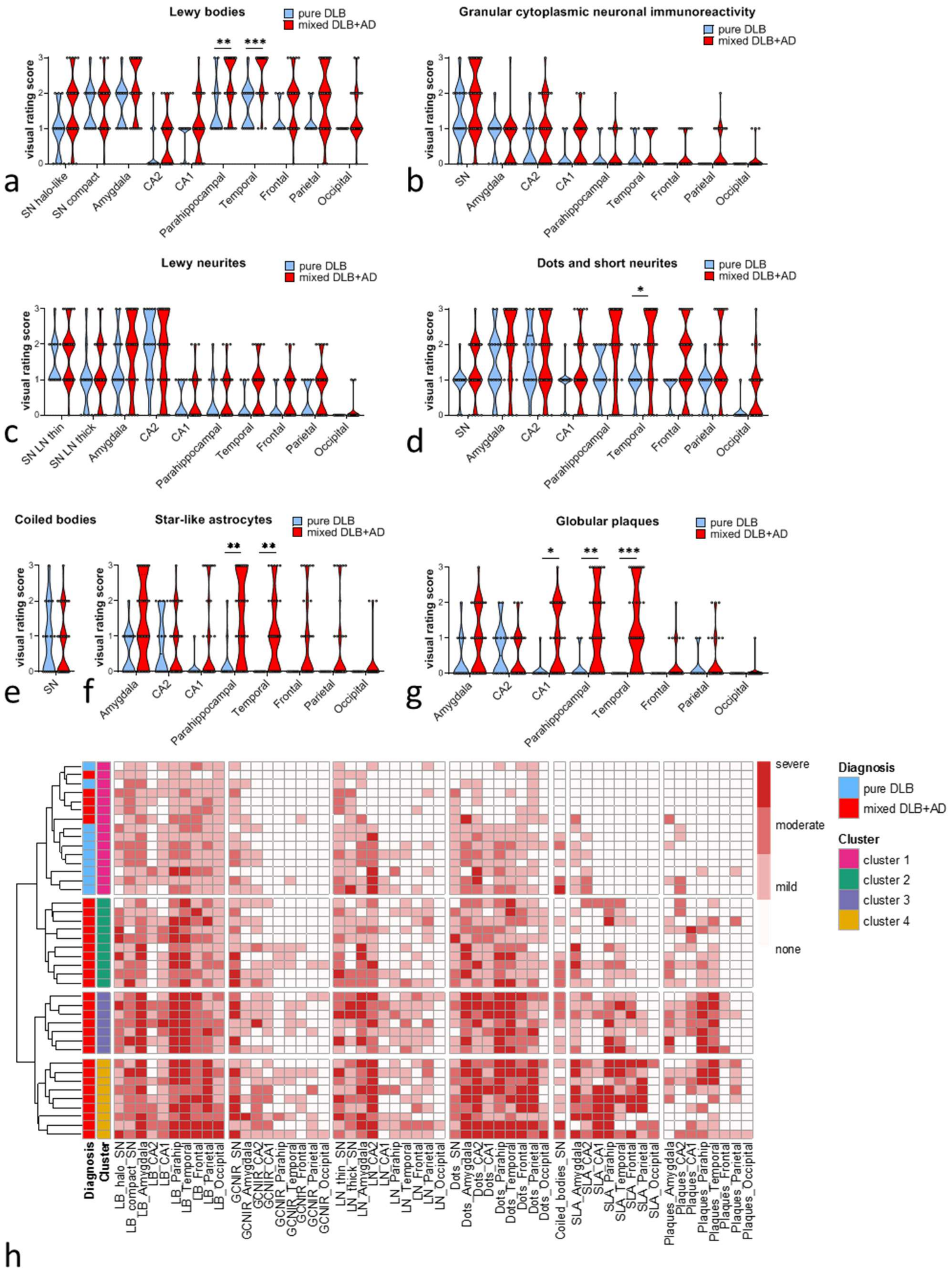

**Supplementary Figure S3. Morphology of αSyn positivity in pure DLB compared to mixed DLB+AD.** In mixed DLB+AD, **a**) Lewy bodies were more abundant in the parahippocampal and temporal cortex, **b**) granular cytoplasmic neuronal immunoreactivity and **c**) Lewy neurites were not increased in specific regions, **d**) dots and short neurites were more abundant in the temporal cortex, **e**) coiled bodies in the substantia nigra were not increased, **f**) star-like astrocytic pathology is more abundant in the parahippocampal and temporal cortex and **g**) globular plaques are more abundant in the CA1 region, parahippocampal and temporal cortex. **h**) Hierarchical cluster analysis of DLB brain donors with a complete dataset (n = 41). All pure DLB donors cluster together in Cluster 1 based on the visual scores of different αSyn morphologies over the brain. These donors have less αSyn-immunoreactive Lewy bodies, dots, star-like astrocytes and plaques than donors in Clusters 2, 3 and 4.

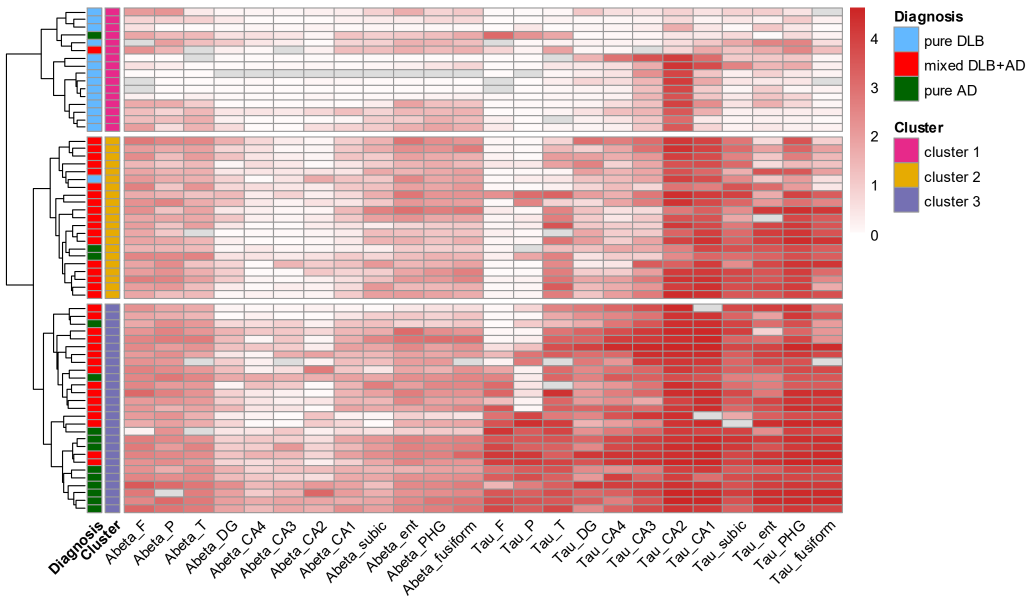


**Supplementary Figure S4. Cluster analysis based on log-transformed amyloid-β and p-tau load in all cases.** Cluster 1 contained 14 out of 15 pure DLB cases. Amyloid-β and p-tau pathology was low in the pure DLB cases, except for p-tau pathology in the CA2 region in a subset of the pure DLB cases. Cluster 2 contained mostly mixed DLB+AD cases, whereas cluster 3b contained mostly pure AD cases and showed the highest levels of amyloid-β and p-tau pathology.

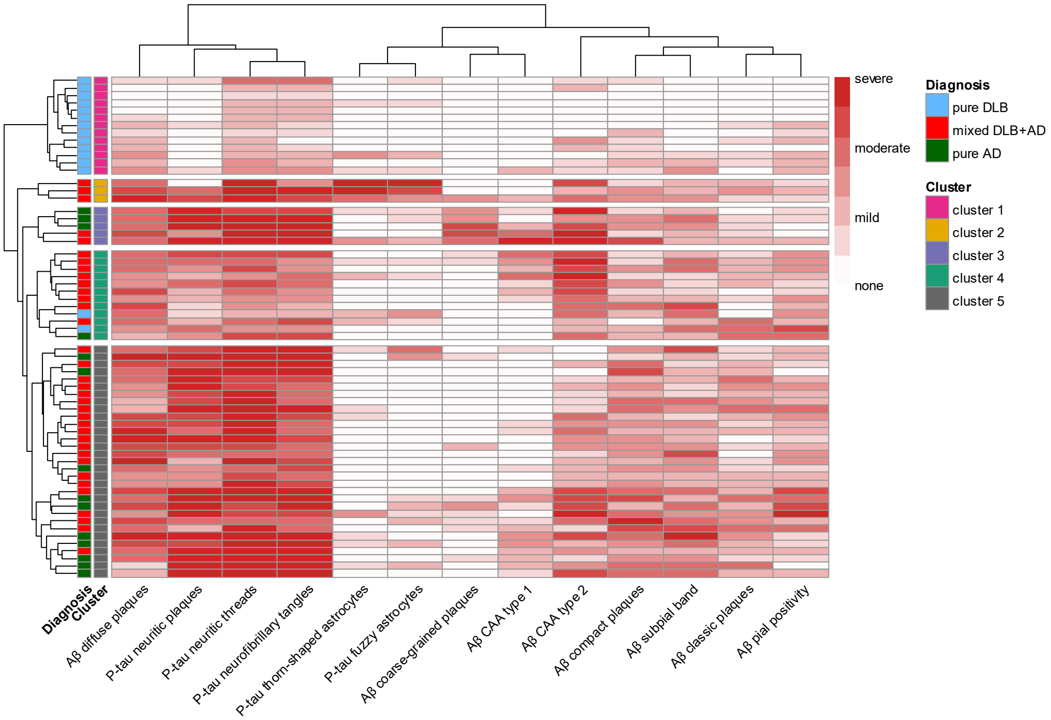


**Supplementary Fig S5. Cluster analysis of cases based on morphology of amyloid-β and p-tau positive structures reveals similar morphology in mixed DLB+AD and pure AD.** Based on the mean visual rating scores of neocortical amyloid-β and tau, 13 out of 15 pure DLB donors clustered in Cluster 1. In the subsequent clusters, the mixed DLB+AD donors were further separated, showing a larger heterogeneity within the mixed DLB+AD group than within the pure DLB group. In Cluster 2, three mixed DLB+AD donors with abundant ARTAG clustered. Cluster 3 contained two mixed DLB+AD donors with abundant coarse-grained plaques and CAA type 1. The other mixed DLB+AD donors could be further separated into two clusters: Cluster 4 with relatively little AD-related p-tau pathology compared to other mixed DLB+AD donors and a high frequency of CAA type 2 pathology, and Cluster 5 with relatively much AD-related p-tau pathology. whereas mixed DLB+AD and pure AD cases clustered together. CAA = cerebral amyloid angiopathy.

1. Ruifrok, A.C. and D.A. Johnston, *Quantification of histochemical staining by color deconvolution.* Analytical and Quantitative Cytology and Histology, 2001. **23**(4): p. 291-9.
